# Beware of Data Leakage from Protein LLM Pretraining

**DOI:** 10.1101/2024.07.23.604678

**Authors:** Leon Hermann, Tobias Fiedler, Hoang An Nguyen, Melania Nowicka, Jakub M. Bartoszewicz

**Affiliations:** Hasso Plattner Institute, Digital Engineering Faculty, University of Potsdam; Computer Science and Artificial Intelligence Laboratory, Massachusetts Institute of Technology

**Author notes:** Equal contribution; corresponding authors. Equal contribution (Leon Hermann, Tobias Fiedler, Hoang An Nguyen). {, }.

**Keywords:** Data Leakage, Protein Representation Learning, Protein Property Prediction, Protein Language Models, Thermostability Prediction

## Abstract

Pretrained protein language models are becoming increasingly popular as a backbone for protein property inference tasks such as structure prediction or function annotation, accelerating biological research. However, related research oftentimes does not consider the effects of data leakage from pretraining on the actual downstream task, resulting in potentially unrealistic performance estimates. Reported generalization might not necessarily be reproducible for proteins highly dissimilar from the pretraining set. In this work, we measure the effects of data leakage from protein language model pretraining in the domain of protein thermostability prediction. Specifically, we compare two different dataset split strategies: a pretraining-aware split, designed to avoid similarity between pretraining data and the held-out test sets, and a commonly-used naive split, relying on clustering the training data for a downstream task without taking the pretraining data into account. Our experiments suggest that data leakage from language model pretraining shows consistent effects on melting point prediction across all experiments, distorting the measured performance. The source code and our dataset splits are available at https://github.com/tfiedlerdev/pretraining-aware-hotprot.

## 1 Introduction

Recent success of transformer-based large language models in Natural Language Processing (NLP) such as BERT [4] and GPT4 [26] has inspired the development of protein language models (PLMs) that can learn powerful representations of protein sequences. Models such as ESM2 [19] and ProtT5 [6] have been pretrained in a self-supervised fashion via the masked language modeling objective on vast numbers of protein sequences. They have been shown to create meaningful protein representations for use in downstream tasks, such as protein structure prediction and function annotation [19]. Autoregressive PLMs are in turn capable of both conditional and unconditional protein sequence generation, including finetuning for target properties [21, 8, 24, 22, 13, 23].

By enabling these applications, PLMs have become an influential technology for computational protein design and analysis, advancing protein research by facilitating various tasks, e.g., estimating the thermostability of unseen protein sequences [16]. However, utilizing representations generated by PLMs comes with unnoticeable challenges when evaluating a downstream tool’s performance: performance metrics are usually acquired on test data, which is supposed to resemble real-world data samples. While test data should follow the same distribution as the training data, its samples should be as independent from the training data as possible. Otherwise, a model’s capability to ‘remember’ the training samples is evaluated, instead of its capability to generalize to truly unseen data. The impact is particularly high on applications based on PLMs. PLMs are pretrained on a vast collection of protein sequences, potentially incorporating almost every documented protein, of which only a fraction is being held out for testing. If the pretraining split is disregarded when choosing a test set for a downstream application, the test set will certainly contain samples that were already used or samples that are related to proteins in the pretraining set. Thus, the PLM serving as the backbone for the downstream model could potentially produce higher-quality embeddings for proteins seen during pretraining, resulting in unrealistic performance estimates for the downstream model. This is especially relevant as these models are often used to predict properties of previously unknown proteins or to design new protein sequences, which might not have been seen during pretraining [9, 3, 17, 14].

To investigate the significance of pretraining data leakage, we measure its effects in the protein thermostability prediction task. Protein stability is a fundamental property that influences many biological processes, such as enzymatic activity, protein-protein interactions, and protein folding. The melting point of a protein, which is defined as the temperature at which the protein loses its native structure and unfolds, is a key indicator of its stability. Accurate prediction of the melting point of a protein is therefore crucial for assessing the real-world applicability of a protein for use cases in medicine or other areas [18].

We compare pretraining-aware and pretraining-unaware data split strategies to measure the effect of information leakage between the training and test data sets. We run these experiments with multiple sizes of the ESM2-family models, ESM-1b and ESMFold. In each of them, a regression head is attached on top of the aggregated protein sequence embeddings, with the protein language model receiving protein sequences as input and the regression head outputting their melting points. As training and test data we utilize the Meltome Atlas [15], which contains numerous protein sequences with multiple experimentally determined melting point measurements per protein.

Our experiments show that data leakage from pretraining consistently inflates the performance when compared to experiments run on the pretraining-aware dataset split. This shows that pretraining data leakage distorts comparisons of different protein language models when used for melting point prediction and highlights the importance of proper dataset splitting in the world of proteins.

## 2 Related Work

### 2.1 Data Leakage

Data leakage is a known challenge in the prediction of protein-protein interactions [1]. Bushuiev et al. [1] highlight problems that occur in existing splitting strategies for protein datasets. Here, we extend on this issue by showing the data leakage between pretrained protein language models and a downstream task. Ding and Steinhardt [5] show that PLMs like ESM2 are biased by unequal sequence sampling across the tree of life, as in the popular protein databases, e.g. UniProt [30], some species are represented by orders of magnitude more samples than others. Ding and Steinhardt demonstrate this by examining protein design’s effects on thermostability following the design methodology proposed by Zhu et al. They measure the thermostability of proteins from thermophilic species before and after protein redesign, utilizing the protein melting temperature predictor DeepSTABp [16]. The results show a decrease in predicted melting points after the redesign, indicating an overall decrease in thermostability [5]. The problem of data leakage is also explored beyond protein-related research. Whalen et al. [31] describe leaky processing as a major pitfall when working with machine learning in genomics. In their work, they explore several other challenges that are not apparent at first sight (e.g. confounding, distributional differences, etc.) due to the inherent complexities of biological systems. They propose further ideas to reduce the risk of encountering these pitfalls [31].

### 2.2 Protein thermostability prediction

Chen et al. [2] conducted a study to predict protein thermostability using both classification and regression methods. They have implemented different methods in their study, including Factorized Sparse Tuning, Structure-Aware Pretraining with AlphaFold2, and creative approaches like mixup feature augmentation and worst-case feature augmentation. The proteins in their dataset are clustered by temperature ranges at a sequence similarity threshold of 50% by employing the CD-Hit algorithm [10]. Unlike in this study, Chen et al. do not use a pretrained PLM and instead fully fine-tune their models, which reduces their risk of data leakage from a differing pretraining split [2].

DeepSTABp is a thermostability predictor for protein sequences [16]. It employs ProtT5 [6] as a transformer-based protein language model combined with feature extraction methods to predict the melting points of proteins. The dataset used by the authors includes the Meltome Atlas [15] and additional proteins from other studies [11, 29]. However, they do not consider pretraining leakage in their study and instead split their dataset randomly into train and validation sets.

Dallago et al. [3] explore representation learning with protein language models of proteins from various landscapes across different protein functions. Their study results in a benchmark, *FLIP*, that also includes thermostability prediction for protein sequences. Their work does not take the pretraining-related leakage of the utilized PLM into account as well [3].

## 3. Methods

### 3.1 ESM1 & ESM2

Evolutionary Scale Modelling 1 and 2 (ESM1 and ESM2) are transformer-based protein language models that have been pretrained on the masked language modeling (MLM) objective to create representations of proteins that are useful for downstream tasks such as protein structure prediction [20]. Pretraining of ESM2, as well as ESM1, has been performed on a dataset based on UniRef [28]. 10% of UniRef50 protein clusters have been held out from pretraining, acting as an evaluation set. The UniRef50 clusters are found with MMseqs2 [27] with a minimum 50% sequence similarity between proteins in the same cluster. ProtT5 [6] data splits have not been made available, which is why we chose the ESM models for our experiments. We focus on masked language models as common backbones for protein property predictors.

For the task of melting point prediction, a variable size input – the protein sequence – is given, and a fixed size output – the melting point – is the prediction target. As the number of embeddings produced by ESM2 depends on the length of the given protein sequence, an aggregation method to obtain a sequence-wide representation of fixed size is needed. The fixed-size representation can then be passed into a simple fully connected neural network called a regression head to predict the protein’s melting point. To acquire the fixed-size representation, we calculate the mean over the sequence of embeddings.

### 3.2 ESMFold

ESMFold [19] is an ESM2-based model for protein structure prediction. To predict the structure of a protein, ESMFold performs a series of steps. First, the input sequence is passed through ESM-2, the language model that generates a representation sequence. Then, the representations are fed into a series of folding blocks, where each block updates the representations that are subsequently passed into the structure module. Within the folding module, pairwise representations include a latent representation for every pair of amino acids in the corresponding sequence. At the same time, the sequence representation contains a latent representation for every amino acid in the corresponding sequence. ESMFold uses both kinds of representations for protein structure prediction, but fully paired representations use considerable amounts of memory. Therefore, we investigated whether the sequence representations updated by the folding modules could contain additional information useful for thermostability prediction, compared to standard ESM2 embeddings.

### 3.3 Dataset

The training dataset utilized in this study is the Meltome Atlas, which covers a total of 201,282 samples consisting of 34,672 unique protein sequences from 13 different organisms. For our experiments, we removed proteins longer than 3000 amino acids to avoid exceeding hardware constraints. The average number of thermostability measurements per protein is 5.81 [15]. The experiments were conducted on two different splits. The first split, *FLIP*, was introduced by Dallago et al. [3]. The authors employed a clustering strategy using MMseqs2 with a sequence similarity of 20%, which produced clusters of proteins with at least 20% similarity to the seed protein of the cluster, i.e. its longest sequence. Their split does not consider the ESM pretraining split and instead randomly samples all protein clusters to create a train, validation, and test set. Consequently, samples from the ESM pretraining set can be found in the *FLIP* test set, making it prone to data leakage from PLM pretraining. Across the sets, 145 proteins were removed due to them exceeding a length of 3000. In total, the dataset consists of 22,335 samples for the training set, 2,466 for the validation set, and 3,115 for the test set. The training and validation sets contain 18,349 and 2,362 unique proteins respectively, while the test set only contains a single measurement per protein.

The second split avoids the risk of data leakage from PLM pretraining. We call it ESM Pretraining-Aware split (*EPA*). Unlike the *FLIP* split, *EPA* employs the UniRef50 clustering to make it compatible with the ESM pretraining clustering. Consequently, clusters here include proteins with at least 50% similarity to the cluster’s seed protein. Due to the increased sequence similarity threshold, the clusters contain fewer proteins, and more clusters are formed. Consequently, proteins within a cluster have a more similar protein sequence. For example, two proteins with 40% sequence similarity will be in separate clusters in the *EPA* split but in the same cluster for the *FLIP* split.

As described in Section 3.1, certain UniRef50 clusters have been excluded from the ESM pretraining data set, acting as the ESM evaluation set. In the *EPA* split, we make use of this by including only those samples that are part of the ESM evaluation clusters in the evaluation set. Therefore, proteins of the same cluster as proteins seen during ESM pretraining cannot be part of our evaluation set. We further assign all samples belonging to randomly selected two-thirds of the held-out protein clusters as the validation set and the remaining one-third as the test set. All samples belonging to other UniRef50 clusters are selected for training. We select the median of all measurements for a given protein sequence, resulting in one measurement per protein sequence. Across the sets, 169 proteins were removed due to them exceeding a length of 3000. In total, the training, validation, and test sets contain 30,844, 2,113, and 1,082 respectively, summing up to a total of 34,208 samples.

## 4 Experiments

We conducted the experiments by either transfer learning the frozen protein embeddings or finetuning the backbone models in addition to the thermostability predictor. The finetuning experiments aimed to increase the quality of the generated embeddings to improve the thermostability prediction capabilities of the following predictor model. Because of the associated computational cost, we only run finetuning on the 35M parameter variant of ESM2. All experiments were replicated for the dataset splits:

1. *EPA*: ESM Pretraining-Aware split of thermostability measurements
2. *FLIP*: Split pattern as defined in the *FLIP* paper (not taking ESM pretraining into account)

The embeddings were mean-aggregated along the protein sequence length dimension, before being passed into the thermostability predictor model. The pretrained backbone models can be found at the ESM GitHub repository [7]. For all of the experiments, we used the regression head for mean aggregation as described by Dallago et al. [3], without conducting a further hyperparameter search. It is a simple, two-layer, fully connected network that maps from the embedding vector to a same-sized hidden representation, followed by ReLU activation and subsequently mapped to the scalar melting point. All experiments used mean squared error as their loss function during training and were trained for 30 epochs. We measured the models performance on the validation dataset after each epoch and the model state was saved. After training the best-performing model checkpoint was chosen and validated on the test set. The experiments were conducted by training the thermostability predictor model on protein embeddings of multiple model backbones:

1. ESM-1b (650M parameters), frozen
2. ESM2 (35M parameters), frozen & finetuned
3. ESM2 (650M parameters), frozen
4. ESM2 (3B parameters), frozen
5. ESM2 (15B parameters), frozen
6. ESMFold sequence representations (utilizing ESM2-650M), frozen

Varying ESM2 model sizes were examined to measure the broad impact of data leakage independent of the underlying backbone model size. As ESM-1b was already applied to the thermostability prediction task in the *FLIP* study [3] and ESM-1b was trained with the same dataset split as ESM2, we also included a reproduction of the *FLIP* findings using their exact pipeline as an additional comparison. As embeddings created with ESMFold contain structural information that should help in predicting thermostability, these embeddings were also included in the experiments.

## 5 Results

The *EPA* split reduces data leakage from PLM pretraining but employs a clustering strategy with a sequence similarity threshold of 50%. This allows proteins with e.g. 40% similarity to be part of the training and the test set, as they do not belong to the same cluster. In contrast, in the *FLIP* split, these proteins would only be represented in either training, validation, or test set. Consequently, assuming that data leakage does not have a significant effect, using the *EPA* split could result in obtaining higher performance estimates compared to the *FLIP* split. Additionally, Figure 2 shows that the *FLIP* test set contains visibly more outliers, e.g., proteins with unusually high melting points, while the train sets of both splits are similarly distributed, not resembling the density of outliers found in the *FLIP* test set. This contributes to the assumption that the *EPA* split should lead to observing higher values of the performance metrics, as long as no detectable data leakage is present. A histogram visualizing this disparity in distributional outliers can be found in Figure A1.

**Fig. 1:**
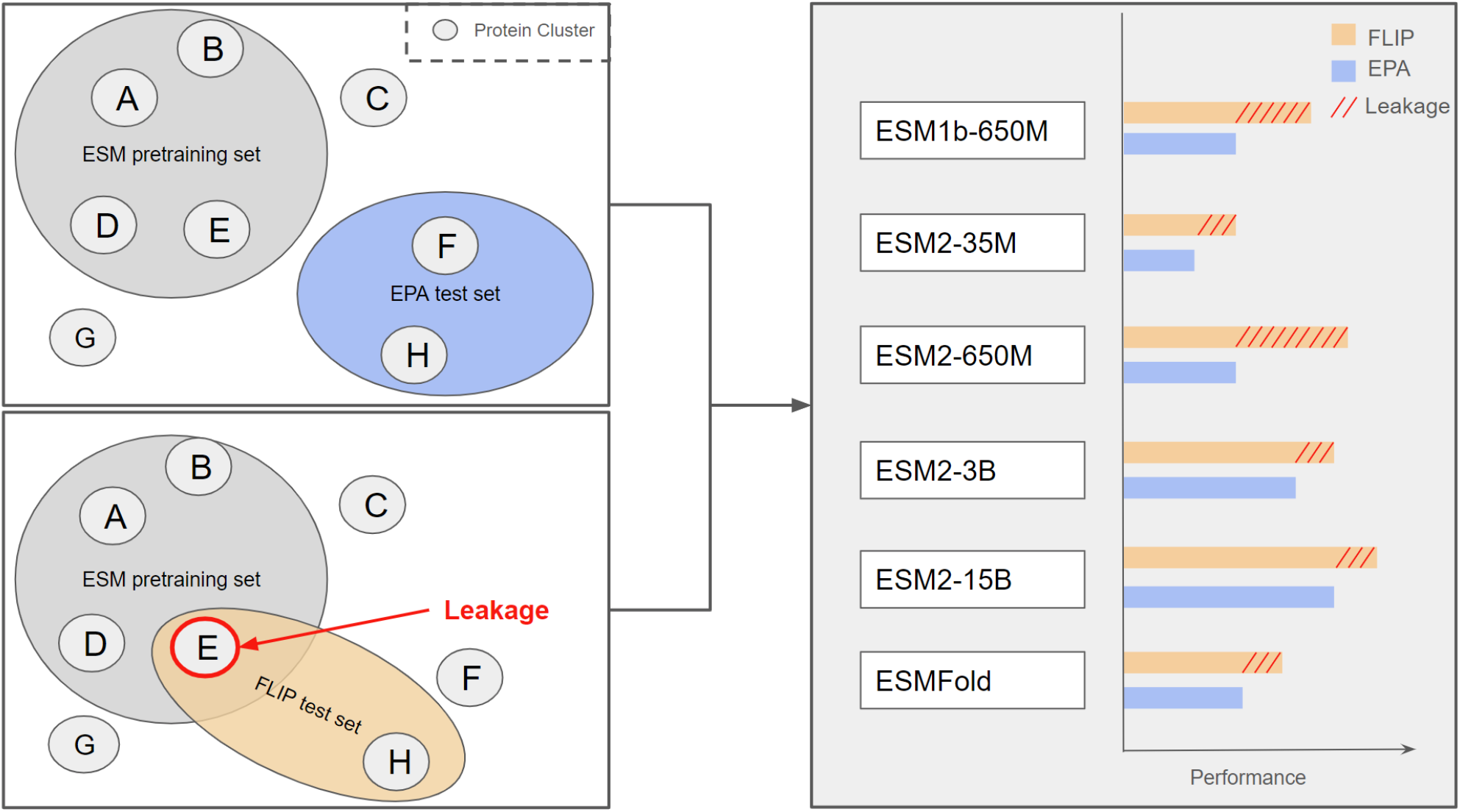
Conceptual drawing of pretraining-aware splitting (*EPA*) vs. pretraining-unaware splitting (*FLIP*). Protein clusters are shown as representatives for all existing UniRef50 protein clusters. A subset of the clusters is used in ESM pretraining. *EPA* takes this into account, while *FLIP* does not, resulting in an overlap in pretraining and downstream task evaluation clusters as highlighted in orange. Protein clusters not assigned to any set are either in ESM pretraining held out test set or not considered for pretraining or the downstream task at all. We conduct multiple experiments on different ESM models to investigate data leakage. Leakage may be detected as unexpectedly inflated performance on the *FLIP* split that cannot be attributed to other sources of split ‘difficulty’, such as sequence similarity thresholds used to remove sequence redundancy.

**Fig. 2:**
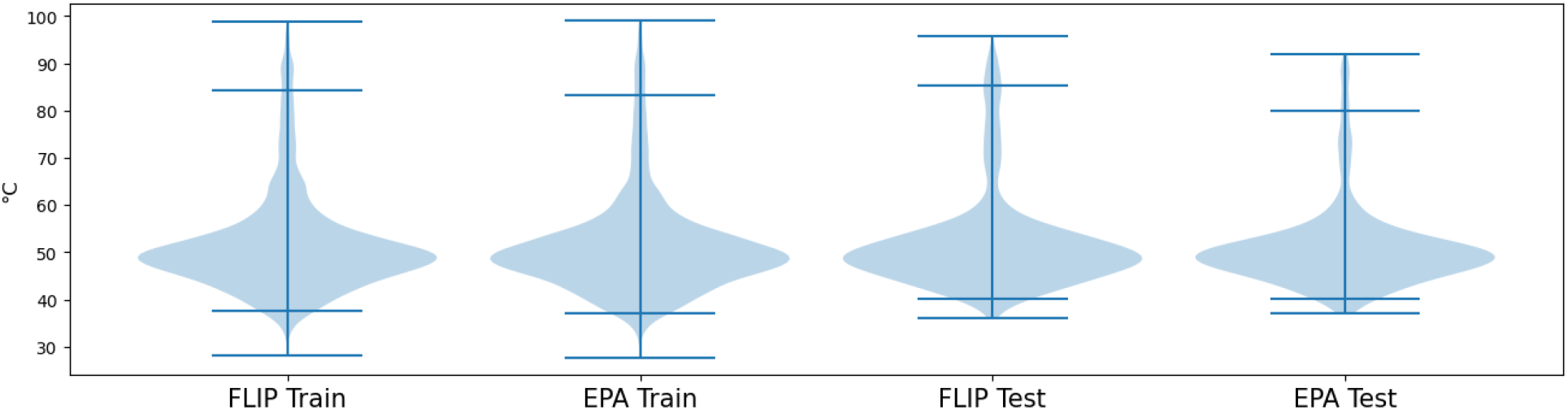
Data distribution of melting points across the utilized splits with 95% quantile range displayed by inner whiskers and extremes marked by outer whiskers. Both train and test sets visibly have a similar distribution. The test sets do differ in terms of upper outliers. Specifically, the test set of the *FLIP* split contains visibly more proteins with higher melting points than the test set of the *EPA* split as well as the equally distributed train sets, making the *FLIP* split more challenging, when not considering any data leakage effects.

To benchmark the model performance, we employed Spearman Rank Correlation Coefficients (SRCC) of prediction and label observed on the test set for each experiment. Ranking-based metrics like Spearman rank correlation are a common choice in protein optimization [3, 25] and ranking-based losses have been shown to improve prediction quality across different protein fitness landscapes [12]. Each experiment was evaluated on the model checkpoint at the epoch with the best validation loss during the training process. We summarize the results in Table 1. The diverse protein embeddings utilized for training are shown alongside the corresponding parameter counts of the underlying models.

**Table 1:** Spearman rank correlation for melting point prediction on the test set of ESM-pretraining-aware split (*EPA*) and *FLIP* split using different PLM backbones. Cells marked with * are related to experiments that we run with the pipeline from the *FLIP* paper.

| Model | EPA | FLIP | Difference |
| --- | --- | --- | --- |
| ESM-1b (650M), frozen | 0.620* | 0.680* | +0.060 |
| ESM2 (35M), frozen | 0.497 | 0.610 | +0.113 |
| ESM2 (650M), frozen | 0.523 | 0.618 | +0.095 |
| ESM2 (3B), frozen | 0.674 | 0.699 | +0.025 |
| ESM2 (15B), frozen | 0.670 | 0.721 | +0.051 |
| ESM2 (35M), finetuned | 0.436 | 0.478 | +0.042 |
| ESMFold seq. rep. (650M), frozen | 0.416 | 0.546 | +0.130 |
| Mean over ESM variants | $\sim 0.548$ | $\sim 0.603$ | +0.055 |

All tested PLM backbone models achieve a higher Spearman rank correlation if they are trained and evaluated on the *FLIP* split compared to the *EPA* split. The Spearman rank correlation on the *EPA* split is statistically significantly worse than on the *FLIP* split (Wilcoxon signed-rank test, *p* = 0.007).

The average Spearman rank correlation for these models on the *EPA* and *FLIP* splits are 0.548 and 0.603 respectively. Additionally, the results show a trend of improved Spearman rank correlation with model size. However, on the *EPA* split, we observe no significant change from the 3B model variant to the 15B variant. On the *FLIP* split, the performance continues to improve. Additionally, the largest differences in Spearman rank correlation are seen for smaller ESM2 variants of up to 650M parameters with the exception of the finetuned 35M parameter model. Our hypothesis that the ESMFold embeddings contain structural information that would increase the performance of the thermostability predictor was not confirmed as the model performed worse compared to similarly sized ESM2 and ESM-1b backbones. Moreover, this model exhibits the largest increase in performance when comparing the *FLIP* split to the *EPA* split.

Prediction residuals of the experiments utilizing the ESM2 15B backbone are visualized in Figure 3. For comparison, two hypothetical models predicting only the average train target for *FLIP* and *EPA* respectively are shown and are labeled “FLIP average” and “EPA average”. We observe a slightly larger 95% quantile range, indicated by the inner whiskers, for both *FLIP* plots compared to the *EPA* split. The increased residual range of the “FLIP average” plot compared to the “EPA average” plot indicates that the *FLIP* split is indeed the more challenging one, disregarding data leakage effects.

**Fig. 3:**
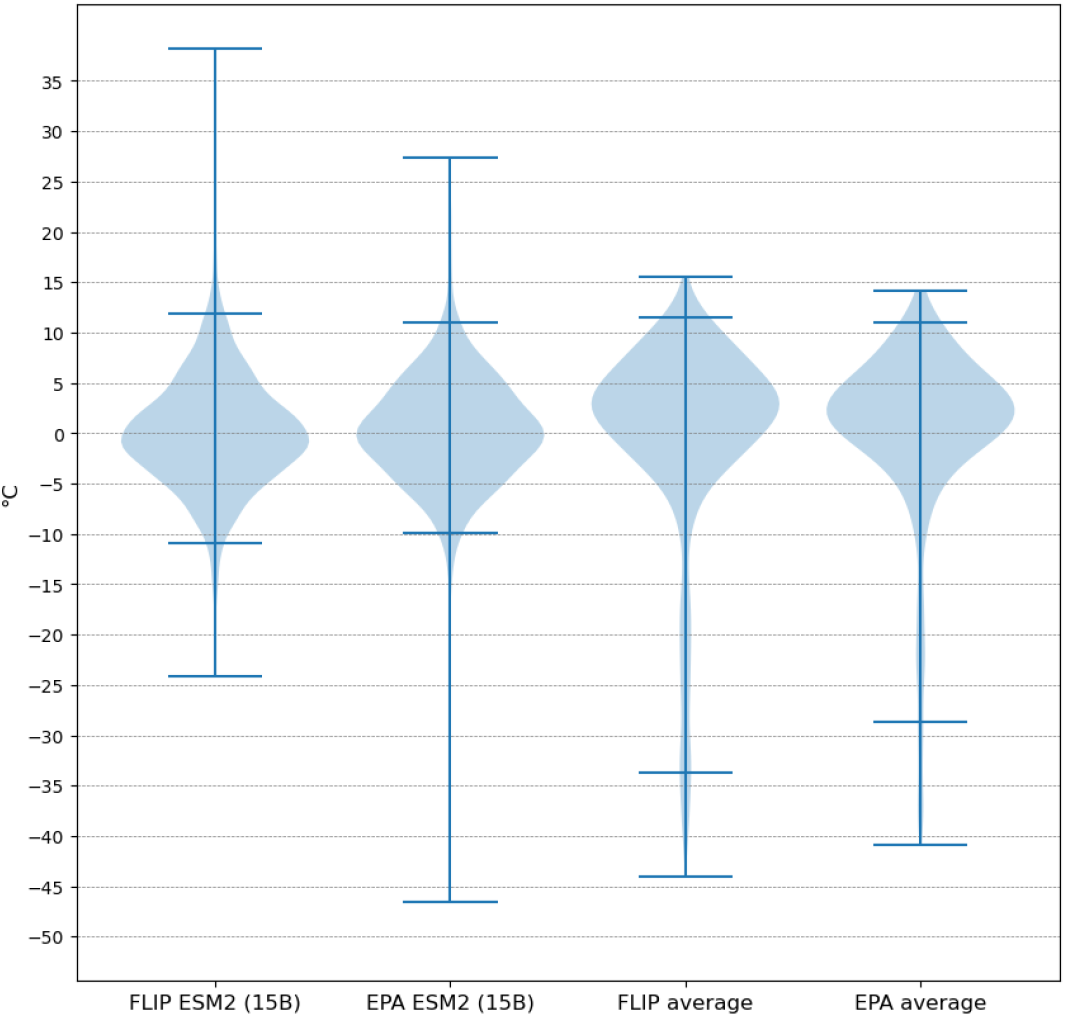
Prediction residuals of the model trained using the ESM-2 15B backbone compared to a naïve average predictor, i.e., a model that just predicts the respective training set’s average melting point for all samples. The 95% quantile range is displayed by inner whiskers and extremes marked by outer whiskers. The smaller 95% quantile ranges and zero-centrality of the residuals of the trained models clearly show that they indeed do out-perform the naive predictor.

## 6 Discussion

Our study examines the extent and effects of pretraining data leakage within the protein language model domain. The results show that all models achieved higher performance on the data leakage-prone *FLIP* split compared to the data leakage-free *EPA* split.

Our analysis indicates that data leakage is the primary factor contributing to these observations. Upon further examination of Figure 2 it becomes evident that the test set of the *FLIP* split contains a greater number of proteins with higher melting points compared to the test set of the *EPA* split. This makes the test set of the *FLIP* split more challenging for the thermostability prediction task, due to the limited availability of very thermostable proteins in the training sets of both splits. Furthermore, the *FLIP* split employs a different clustering strategy with 20% protein sequence identity instead of 50% as implemented in the UniRef50 clustering. Therefore, protein sequences in the *FLIP* test set share less similarity with protein sequences from the train set, making it even more difficult to generalize on the unseen test set. Despite these facts, the models achieve a higher performance on the *FLIP* split. This inconsistency can be explained by the data leakage that comes from the disregard of clustering and splitting of the datasets used during pretraining. The difference between *FLIP* and *EPA* is especially evident for models below 3B parameters. A potential cause for this could be that smaller models do not generalize well beyond their training data for a complex task such as protein language modeling. Since data leakage is present in the *FLIP* split, potential overfitting to the training data becomes visible in the test metrics.

As we have only measured the effects of pretraining data leakage in the protein thermostability prediction task, the applicability of these findings to other tasks needs to be confirmed. However, as ESM2 is not specifically designed for the thermostability prediction task, but rather to learn protein representations that capture properties beyond the amino-acid sequence, we believe that the effects of pretraining data leakage apply to other tasks as well. Furthermore, a limitation of this study is the exclusive testing on ESM backbone models, which were trained on the masked language modeling objective. While we expect other PLMs with the same training objective to exhibit similar effects of pretraining data leakage, future work should expand on other models and models that were trained on different objectives. Additionally, the experiments were conducted on only two different splits. This may indicate that the observed effect could be caused by differences in sampling the train, validation, and test sets or other unknown effects. To exclude the possibility of such factors, we ran several experiments utilizing different model backbones and we employed a Wilcoxon test to confirm whether these findings are statistically significant. The test yielded a p-value of 0.007 which we consider significant. To further eliminate random factors, reproducing these experiments with multiple data leakage-prone splits could validate our findings to a greater extent.

## 7 Conclusion

Data leakage from pretraining is not always considered in protein language model-related research. In our work, we have shown that it has a significant effect on the measured performance to an extent that could be a major limitation to the conclusions drawn from commonly used benchmarking procedures. Although this study is limited to a single regression task and a family of related pretrained models, we expect the conclusions to apply also beyond this particular setup. Therefore, we call the researchers using protein language models to consider potential data leakage from pretraining and carefully choose their data splitting strategy to guarantee the validity and reproducibility of the produced results.

## Code and Data Availability

The source code and our dataset splits are available at https://github.com/tfiedlerdev/pretraining-aware-hotprot.

## Acknowledgements

This work was partially supported by the HPI-MIT Research Program “Designing for Sustainability” funded by the Hasso Plattner Foundation. We thank Bernhard Renard for helpful discussions and comments.

## A Appendix

**Fig. A1:**
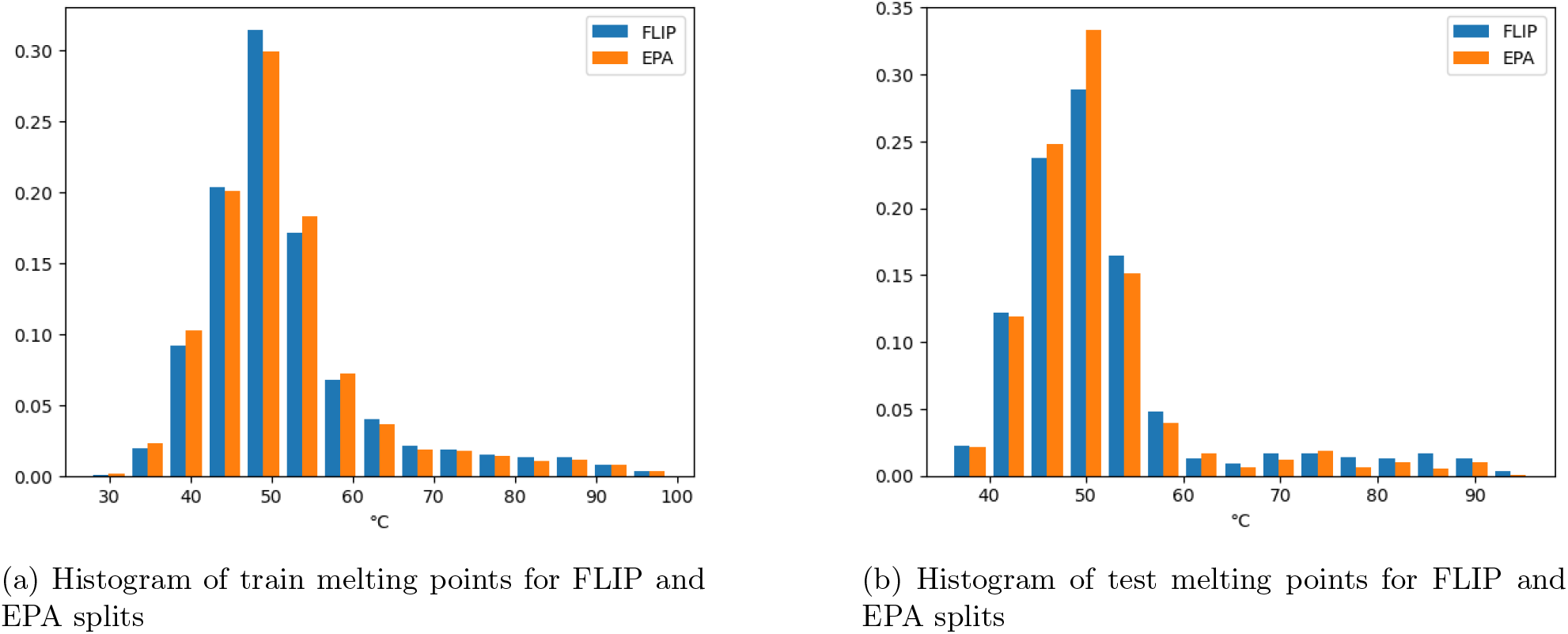
Data distribution of melting points across different splits. Both train and test sets visibly have a similar distribution. The test sets do differ in terms of upper outliers though. Specifically, the test set of the FLIP split contains visibly more proteins with higher melting points (i.e. greater than 75°C) than the test set of the EPA split as well as the more similarly distributed train sets, making the FLIP split more challenging, when not considering any data leakage effects.

## References

[1] Anton Bushuiev et al. Revealing data leakage in protein interaction benchmarks. 2024. doi: 10.48550/ARXIV.2404.10457. url: https://arxiv.org/abs/2404.10457.

[2] Tianlong Chen et al. “Hotprotein: A novel framework for protein thermostability prediction and editing”. In: The Eleventh International Conference on Learning Representations. 2022.

[3] Christian Dallago et al. “FLIP: Benchmark tasks in fitness landscape inference for proteins”. In: bioRxiv (2021). doi: 10.1101/2021.11.09.467890. eprint: https://www.biorxiv.org/content/early/2021/11/11/2021.11.09.467890.full.pdf. url: https://www.biorxiv.org/content/early/2021/11/11/2021.11.09.467890.

[4] Jacob Devlin et al. BERT: Pre-training of Deep Bidirectional Transformers for Language Understanding. 2019. 1810.04805[cs.CL].

[5] Frances Ding and Jacob Steinhardt. “Protein language models are biased by unequal sequence sampling across the tree of life”. In: (Mar. 2024). doi: 10.1101/2024.03.07.584001. url: http://dx.doi.org/10.1101/2024.03.07.584001.

[6] Ahmed Elnaggar et al. ProtTrans: Towards Cracking the Language of Life’s Code Through Self-Supervised Deep Learning and High Performance Computing. 2020. doi: 10.48550/ARXIV.2007.06225. url: https://arxiv.org/abs/2007.06225.

[7] Meta Facebookresearch. Facebookresearch/ESM: Evolutionary scale modeling (ESM): Pretrained language models for proteins. url: https://github.com/facebookresearch/esm.

[8] Noelia Ferruz, Steffen Schmidt, and Birte Höcker. “ProtGPT2 is a deep unsupervised language model for protein design”. In: Nature communications 13.1 (2022), p. 4348.

[9] Zachary N Flamholz, Steven J Biller, and Libusha Kelly. “Large language models improve annotation of prokaryotic viral proteins”. In: Nature Microbiology (2024), pp. 1–13.

[10] Limin Fu et al. “CD-HIT: accelerated for clustering the next-generation sequencing data”. In: Bioinformatics 28.23 (Oct. 2012), pp. 3150–3152. issn: 1367-4811. doi: 10.1093/bioinformatics/bts565. url: http://dx.doi.org/10.1093/bioinformatics/bts565.

[11] Carina Groh et al. “Mitochondrial dysfunction rapidly modulates the abundance and thermal stability of cellular proteins”. In: Life Science Alliance 6.6 (Mar. 2023), e202201805. issn: 2575-1077. doi: 10.26508/lsa.202201805. url: http://dx.doi.org/10.26508/lsa.202201805.

[12] Alex Hawkins-Hooker, Paul Duckworth, and Oliver Bent. “Preferential Bayesian Optimisation for Protein Design with Ranking-Based Fitness Predictors”. In: ().

[13] Daniel Hesslow et al. “Rita: a study on scaling up generative protein sequence models”. In: arXiv preprint 2205.05789 (2022).

[14] C Hsu et al. “Learning inverse folding from millions of predicted structures. bioRxiv 2022”. In: preprint (2022).

[15] Anna Jarzab et al. “Meltome atlas—thermal proteome stability across the tree of life”. In: Nature methods 17.5 (2020), pp. 495–503.

[16] Felix Jung et al. “DeepSTABp: A Deep Learning Approach for the Prediction of Thermal Protein Stability”. In: International Journal of Molecular Sciences 24.8 (Apr. 2023), p. 7444. issn: 1422-0067. doi: 10.3390/ijms24087444. url: http://dx.doi.org/10.3390/ijms24087444.

[17] Henry R Kilgore et al. “Protein codes promote selective subcellular compartmentalization”. In: bioRxiv (2024), pp. 2024–04.

[18] Benjamin Kumwenda et al. “Analysis of Protein Thermostability Enhancing Factors in Industrially Important Thermus Bacteria Species”. In: Evolutionary bioinformatics online 9 (Aug. 2013), pp. 327– 42. doi: 10.4137/EBO.S12539.

[19] Zeming Lin et al. “Evolutionary-scale prediction of atomic level protein structure with a language model”. In: bioRxiv (2022). bioRxiv 2022.07.20.500902. url: 10.1101/2022.07.20.500902.

[20] Zeming Lin et al. “Language models of protein sequences at the scale of evolution enable accurate structure prediction”. In: bioRxiv (2022). doi: 10.1101/2022.07.20.500902. eprint: https://www.biorxiv.org/content/early/2022/07/21/2022.07.20.500902.full.pdf. url: https://www.biorxiv.org/content/early/2022/07/21/2022.07.20.500902.

[21] Ali Madani et al. “Large language models generate functional protein sequences across diverse families”. In: Nature Biotechnology 41.8 (2023), pp. 1099–1106.

[22] Geraldene Munsamy et al. “Conditional language models enable the efficient design of proficient enzymes”. In: bioRxiv (2024), pp. 2024–05.

[23] Andrea Nathansen et al. “Evaluating Tuning Strategies for Sequence Generation with Protein Language Models”. In: Proceedings of the 18th Machine Learning in Computational Biology meeting. Ed. by David A. Knowles and Sara Mostafavi. Vol. 240. Proceedings of Machine Learning Research. PMLR, 2024, pp. 76–89. url: https://proceedings.mlr.press/v240/nathansen24a.html.

[24] Erik Nijkamp et al. “Progen2: exploring the boundaries of protein language models”. In: Cell systems 14.11 (2023), pp. 968–978.

[25] Pascal Notin et al. “ProteinNPT: improving protein property prediction and design with non-parametric transformers”. In: Advances in Neural Information Processing Systems 36 (2023), pp. 33529–33563.

[26] OpenAI. GPT-4 Technical Report. 2023. 2303.08774[cs.CL].

[27] Martin Steinegger and Johannes Söding. “MMseqs2: sensitive protein sequence searching for the analysis of massive data sets”. In: bioRxiv (2017). doi: 10.1101/079681. eprint: https://www.biorxiv.org/content/early/2017/06/07/079681.full.pdf. url: https://www.biorxiv.org/content/early/2017/06/07/079681.

[28] Baris E. Suzek et al. “UniRef clusters: a comprehensive and scalable alternative for improving sequence similarity searches”. In: Bioinformatics 31.6 (2014), pp. 926–932. doi: 10.1093/bioinformatics/btu739. url: https://doi.org/10.1093/bioinformatics/btu739.

[29] Chris Soon Heng Tan et al. “Thermal proximity coaggregation for system-wide profiling of protein complex dynamics in cells”. In: Science 359.6380 (Mar. 2018), pp. 1170–1177. issn: 1095-9203. doi: 10.1126/science.aan0346. url: http://dx.doi.org/10.1126/science.aan0346.

[30] Uniprot Protein Embeddings. url: https://www.uniprot.org/help/embeddings.

[31] Sean Whalen et al. “Navigating the pitfalls of applying machine learning in genomics”. In: Nature Reviews Genetics 23.3 (Nov. 2021), pp. 169–181. issn: 1471-0064. doi: 10.1038/s41576-021-00434-9. url: http://dx.doi.org/10.1038/s41576-021-00434-9.

[32] Danqing Zhu et al. “Optimal trade-off control in machine learning–based library design, with application to adeno-associated virus (AAV) for gene therapy”. In: Science Advances 10.4 (Jan. 2024). issn: 2375-2548. doi: 10.1126/sciadv.adj3786. url: http://dx.doi.org/10.1126/sciadv.adj3786.

